# DeepCORE: An interpretable multi-view deep neural network model to detect co-operative regulatory elements

**DOI:** 10.1101/2023.04.19.536807

**Authors:** Pramod Bharadwaj Chandrashekar, Hai Chen, Matthew Lee, Navid Ahmadinejad, Li Liu

## Abstract

Gene transcription is an essential process involved in all aspects of cellular functions with significant impact on biological traits and diseases. This process is tightly regulated by multiple elements that co-operate to jointly modulate the transcription levels of target genes. To decipher the complicated regulatory network, we present a novel multi-view attention-based deep neural network that models the relationship between genetic, epigenetic, and transcriptional patterns and identifies co-operative regulatory elements (COREs). We applied this new method, named DeepCORE, to predict transcriptomes in 25 different cell lines, which outperformed the state-of-the-art algorithms. Furthermore, DeepCORE translates the attention values embedded in the neural network into interpretable information, including locations of putative regulatory elements and their correlations, which collectively implies COREs. These COREs are significantly enriched with known promoters and enhancers. Novel regulatory elements discovered by DeepCORE showed epigenetic signatures consistent with the status of histone modification marks.

## Introduction

Gene transcription displays complicated spatiotemporal patterns that vary between cell types, developmental stages, disease phenotypes, and environmental exposures^1,2^. Such variations are regulated by a set of mechanisms that induce or repress gene transcription as part of a large network^3,4^. Many factors participate in gene transcription regulation, such as genetic alterations^5,6^, epigenetic changes^7,8^, and chromatin structure^9–11^. Deciphering and cataloging these regulatory codes are a grand challenge.

Computational mining of multi-omics data is a promising approach to investigate the mechanisms of gene transcriptional regulation. As early attempts, several models used genetic sequence information such as transcription factor binding sites (TFBS) to predict gene transcription levels.^12–18^ However, relying on genetic sequences that cannot capture tissue-specific information is a major limitation of these models. Epigenetic features, such as histone modification marks (HMM), are introduced to address this issue. DeepChrome^19^ is one of the early deep learning method that modelled the relationship between epigenetic and transcriptional profiles. It retrieves the HMM signals in the ±5kbps region around the transcription start site (TSS) of a gene, uses a convolutional neural network (CNN) to extract local features, and feeds these features to a feedforward neural network (FNN) to predict gene transcription levels. ExPecto^20^ expands the TSS-flanking region to 40kbps and includes ChIP-seq data of hundreds of TFs as input. Although these models reported high accuracy of predicting gene transcription levels, they do not identify regulatory elements (REs) in the genome that are essential to understanding the regulatory mechanisms.

Because deep neural networks (DNN) are often considered a black box, extracting biological meanings from these models can be challenging. Recently, several algorithms have been developed to interpret and visualize patterns learned in DNN^21–23^. DeepChrome summarized the HMM patterns coded in the CNN model, reporting consistency with known active and repressive marks. However, a high-level summary cannot identify and locate REs. Furthermore, epigenetic features highly depend on genetic features. For example, chromatin structure changes involving TFBS will have a larger impact on gene transcription than those outside TFBS. In this study, we present a novel method to address this knowledge gap and model co-operative regulatory elements (COREs).

This new method, named DeepCORE, uses a multi-view architecture to integrate genetic and epigenetic profiles in a DNN. It captures short-range and long-range interactions between REs through bidirectional Long short-term memory (BiLSTM). It leverages the attention mechanism to enhance the interpretability of the model and the identification of REs. The output of DeepCORE includes prediction of gene transcription level, locations of REs in the genome, and correlations of multiple REs. We applied DeepCORE on 25 cell lines and showed that DeepCORE has significantly higher accuracies than existing state-of-the-art methods. DeepCORE model has good generalizability across cell lines and identifies COREs with high resolution and enrichment of known promoters and enhancers.

## Methods and Materials

### Overall design and data sets

DeepCORE has two components (**Fig. 1**). The first component is a deep neural network (DNN) that predicts transcription level of a gene based on its genetic and epigenetic features. The second component is an interpreter that analyzes the attention matrices in the DNN to discover COREs.

**Figure 1:**
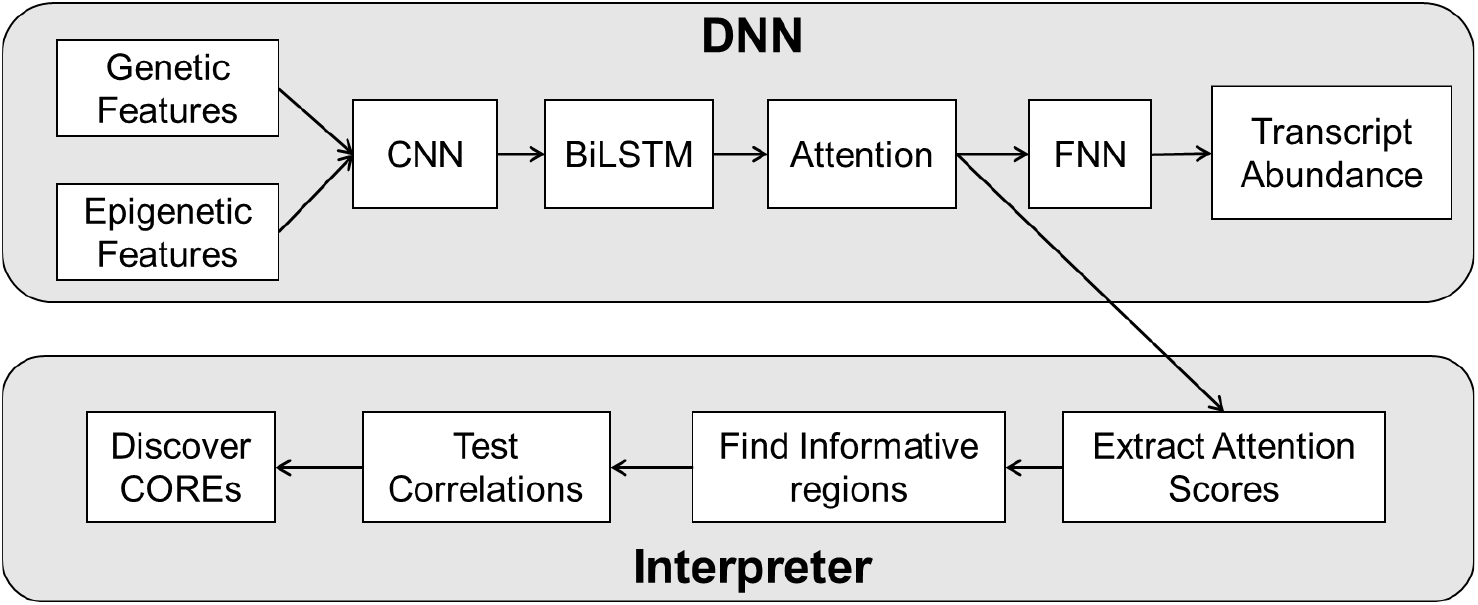
The DeepCORE framework. It consists of two components. In the DNN component, genetic and epigenetic signals go through a CNN layer, a BiLSTM layer, an Attention layer, and a FNN layer to predict transcript abundance of a gene. In the Interpreter component, attention scores extracted from the output of the Attention layer is analyzed to identify informative and correlated regions as COREs.

For a given gene, DeepCORE focuses on the ±5Kbps region surrounding the TSS. To derive genetic features, we extracted the ±5Kbps nucleotide sequences for each gene and converted into a one hot encoding format. This gives us the genetic feature matrix with rows corresponding to the nucleotides and columns corresponding to genomic regions with the value in each cell corresponding to the presence or absence of a specific nucleotide.

To derive epigenetic features, we obtained ChIP-seq data of 25 cell lines from the ENCODE^24^ project and the RoadMap Epigenomics project (REMC)^25^, The ChIP-seq data contained normalized read counts measuring five HMMs including known transcription activator marks (H3K4me, H3K4me3 and H2K27ac) and repressor markers (H3K9me3 and H3K27me3)^26^. Given a gene, we examined the ±5Kbps TSS-flanking region and recorded position-specific normalized read count for each histone modification mark. These data from each cell line were organized into an epigenetic feature matrix with rows corresponding to HMMs and columns corresponding to genomic positions.

The ENCODE and REMC projects also contained RNA-seq data. For each cell line, we obtained the Reads Per Kilobase of transcript per Million mapped reads (RPKM) value for each gene. These data were organized into a single-column vector with rows corresponding to genes.

For each cell line, we removed genes with missing values of RNA-seq data and missing values of ChIP-seq data across all five histone marks.

### Multi-view attention-based DNN

The DNN architecture consists of two separate paths representing the genetic view and the epigenetic view, respectively (**Fig. 2**). Each path starts with a CNN layer consisting of *N*_*CNN*_ filters with size *k* and stride 1. The output of the CNN layer is passed to a ReLU function connected to max pooling over non-overlapping interval of length *p*. These steps produce a vector of size (*N*_*C*_ − *K +* 1)/*p* for filter *f*_*i*_, *i* ∈ {1, …, *N*_*CNN*_} encoding sequence motifs and another vector of equal size encoding combinations of histone modification patterns. To avoid overfitting, dropout^27^ with a probability of 0.5 is applied to the max-pooled vector. While CNN catches various patterns within a genomic region, it does not consider interactions between regions. Since enhancers and promoters that are separated by Kilobase pairs can interact to regulate gene transcription, DeepCORE uses bi-directional long short-term memory (BiLSTM) networks^28^ to capture the short-range and long-range dependencies.

**Figure 2:**
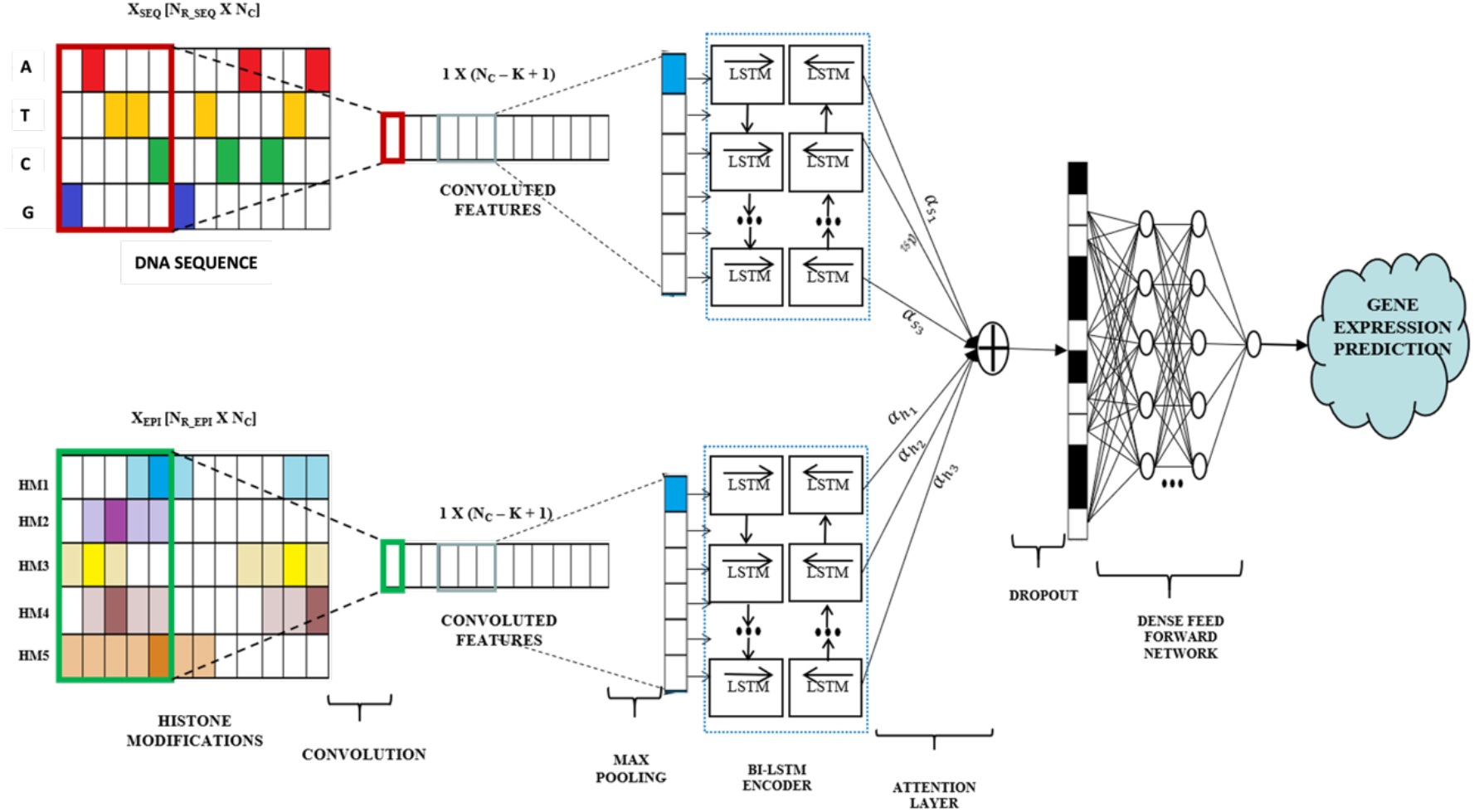
The DNN architecture. The genetic view and the epigenetic view, each consisting of a CNN layer, a BiLSTM layer, and an attention layer, are concatenated before fed into a FNN layer to predict gene transcription level.

As the input sequences to the BiLSTM get longer, it becomes more difficult for the hidden states to capture the context, which reduces the performance^29,30^. To address this issue, DeepCORE employs an encoder-decoder^31^ with attention mechanism^32^. The encoder is the BiLSTM model, and the decoder predicts the importance score of the next genomic region based on importance scores of the regions it has already predicted. This allows the prediction to be made based on a series of important hidden states from the encoder instead of only the last state. Furthermore, DeepCORE highlights the most informative regions contributing to gene transcription regulation by replacing the default softmax function in the attention model with a sparsemax function^33^ that introduces sparsity of probability distribution. The learnt attention is a vector of length equal to the number of output nodes from the CNN layer containing importance score of each genomic region. DeepCORE then joins the two views by concatenating the decoder outputs from each view and giving it to a fully connected FNN to predict continuous gene transcription levels.

### Training DNN

We split the data into disjoint training, validation, and test sets by including 80%, 10%, 10% of all genes via random sampling. The test set was kept hidden and was used only after hyperparameter tuning and parameter learning were completed to avoid information leak. Mean Squared Error (MSE) was computed as the loss function and fed back to the network through backpropagation. We used Adam (Adaptive Moment Estimation) optimizer^34^ to train the model for 100 epochs. Early stopping criteria (the training is stopped if the model performance on the validation set does not improve over 5 epochs) is employed to avoid overfitting.

The optimization was performed at two stages., At the first stage, the hyperparameters in the DNN model were optimized via grid search (**Supplementary Table 1**). The optimal configuration was selected based on the performance on the validation set. The second stage of optimization is done on the attention mechanism for obtaining sparsity. The parameters in the DNN model for both the stages were jointly learned.

### Interpreting attention matrices to discover COREs

For each cell line, we built a multi-view attention-based DNN model. This model contains an attention probability matrix with rows corresponding to genes and columns corresponding to 50bps sliding window (bin) surrounding the TSS. We extract the tissue-specific attention probabilities for the bins for each gene and compute the cumulative distribution (CDF) of this attention probability distribution. We then computed empirical p-values based on the CDF and applied correction of multiple comparison to derive the false discover rate (FDR). Bins with FDR<0.05 indicated genomic regions with significant regulatory function.

After extracting the significant bins for each gene across all tissues, we obtained a matrix with rows representing different tissues and columns representing the bin probabilities. Pearson’s pairwise correlation^35^ was then applied to this matrix that gives us the correlation between different regions in the gene.

Blocks of bins that have significantly correlated attention probabilities and are at least 1kbps apart are putative COREs, i.e., regulatory elements that co-operatively modulate gene transcription.

## Results

### DeepCORE accurately predicts within- and cross-cell line-specific gene transcription levels

We first used two cell lines: E061 (melanocyte cells) and E071(brain hippocampus middle) to assess the performance of DeepCORE DNN model in comparison with two baseline models (single-view DNN using genetic features and epigenetic features alone), and with two state-of-the-art methods (Expecto and DeepChrome). The hyperparameters selected via grid search are K=50, *N*_*CNN*_ =50, p=50, *N*_*LSTM*_ =15, *N*_*ATTN*_ =25, and *N*_*FNN*_ =100. With this setting, genetic sequences and epigenetic signals in each 50bps window are convolved separately. BiLSTM with attention layer produces 200 bins (50bps long), each receiving attention probabilities before being fed to the FNN.

Because the performance of any DNN model is affected by the randomness in weight initialization, we repeated the data set splitting step and the model training step 20 times. Evaluated on the held-out test sets from these 20 repeats, the multi-view DNN reported the lowest error rate in both cell lines (mean and standard deviation of RMSE_*E*061_ = 1.80 ± 0.014 and RMSE_*E*071_ = 1.79 ± 0.015) compared to the two baseline single-view models (genetic-view model: RMSE_*E*061_ = 2.68 ± 0.004 and RMSE_*E*071_ = 2.31 ± 0.002, and epigenetic-view model: (RMSE_*E*061_ = 1.87 ± 0.015 and RMSE_*E*071_ = 1.5 ± 0.055, **Fig. 3A**). It also produced lower error rates than Expecto (RMSE_*E*061_ = 2.86 ± 0.034 and RMSE_*E*071_ = 3.02 ± 0.019). As DeepChrome only outputs binary classification, we converted the continuous prediction scores from DeepCORE and ExPecto using a median cutoff. Again, DeepCORE reported the highest accuracy (F1_*E*061_ = 0.845 ± 0.11 and F1_*E*071_ = 0.749 ± 0.089) compared to the genetic-view model (F1_*E*061_ = 0.69 ± 0.37 and F1_*E*071_ = 0.627 ± 0.062) epigenetic-view model (F1_*E*061_ = 0.834 ± 0.25 and F1_*E*071_ = 0.734 ± 0.084), Expecto (F1_*E*061_ = 0.82 ± 0.057 and F1_*E*071_ = 0.753 ± 0.3), and DeepChrome (F1_*E*061_ = 0.706 ± 0.017 and F1_*E*071_ = 0.69 ± 0.023, **Fig. 3B**).

**Figure 3:**
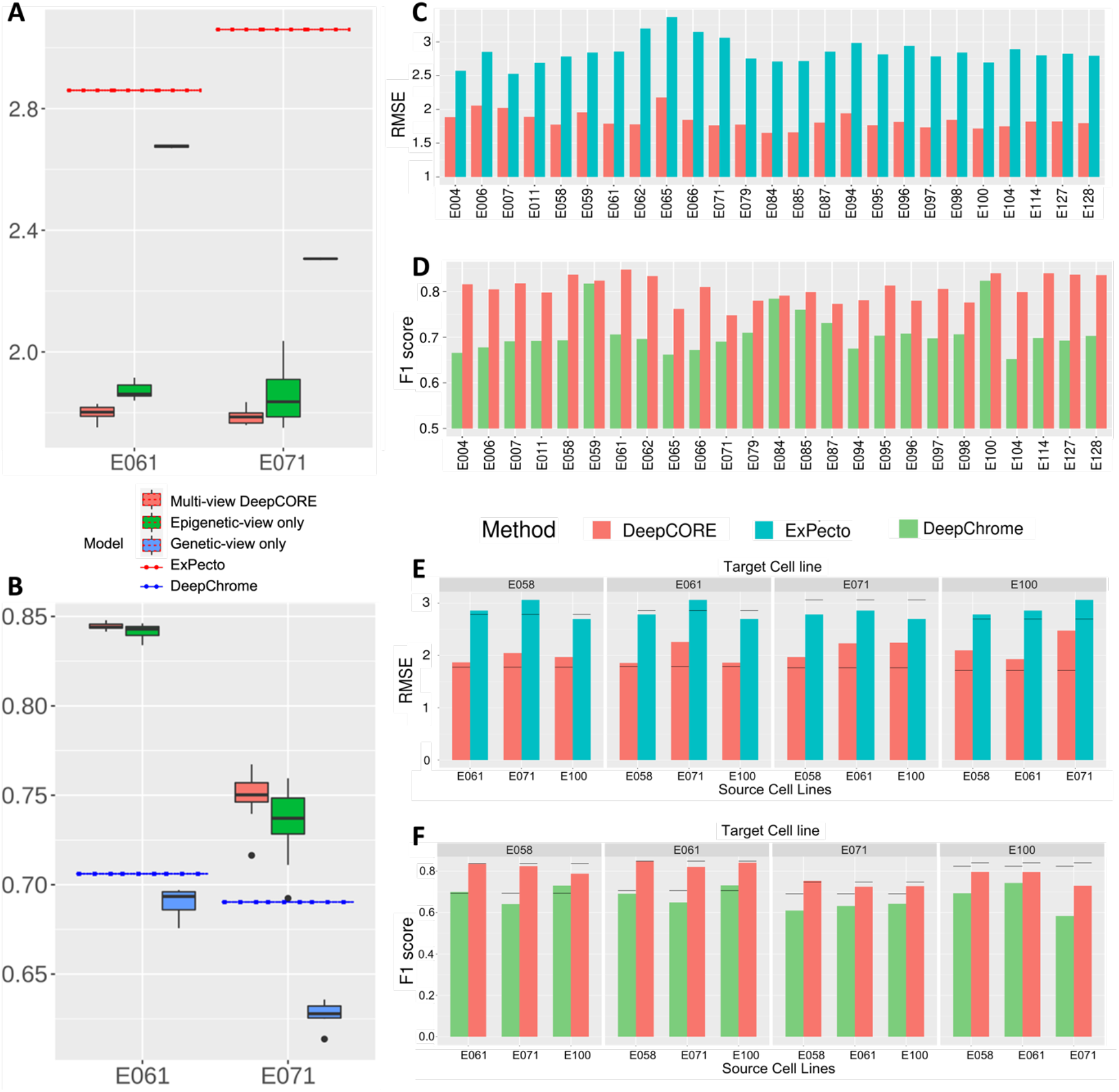
Performance of DeepCORE and other methods. (**A, B**) Evaluated on two cell lines, E061 and E071, the boxplots of RMSE (A) and F1-score (B) show DeepCORE has the lowest error rate and the highest accuracy in predicting gene transcription levels, as compared to single-view DNN, Expecto, and DeepChrome models. (**C, D**) Evaluated on 25 cell lines, DeepCORE has consistently lower error rate than Expecto in predicting continuous gene transcription level (C) and consistently higher accuracy than DeepChrome in predicting binary gene transcription classes (D). (**E, F**) Evaluated on cross-cell-line predictions in which a model trained the source cell line is applied to predict gene transcription in different target cell lines, DeepCORE shows consistently lower error rate than Expecto (E) and higher accuracy than DeepChrome (F). Gray lines denote performance in source cell lines.

We then evaluated the performance of DeepCORE in predicting gene expression levels on 25 cell lines and compared the predictions with Expecto and DeepChrome. Across all cell lines, DeepCORE DNN consistently reported a lower error rate (RMSE) than Expecto (**Fig. 3C)**.The best performance of DeepCORE was observed in the E084 cell line with an RMSE of 1.65, and the lowest performance was observed in the E006 cell line with an RMSE of 2.06. Similarly, DeepCORE consistently reported higher accuracy than DeepChrome on binary classification (**Fig. 3D)**. On average DeepCORE outperformed Expecto and DeepChrome with an improvement of over 10% in most cell lines.

Following the success of DeepCORE DNN on predicting within-cell line gene transcription, we tested if a DNN model trained on one cell line performed well on other cell lines, which indicate the generalizability of the model. We chose four tissues (E058: keratinocyte, E061: melanocyte, E071: brain hippocampus middle, and E100: psoas muscle) which represented very different cell types for this analysis. The results from our analysis indicate that, in general, cross-cell-line predictions have lower performance compared to within-cell-line predictions across all three methods with an exception in the case of Expecto for E071 cell line, where cross-cell-line prediction is better than within-cell-line prediction (**Fig. 3E)**. While the Expecto and DeepChrome showed a huge performance reduction by more than 15% and 10% respectively, the performance of DeepCORE decreased only slightly by 6%. On average, the RMSE error rate of DeepCORE was 27.5% lower than that of Expecto in cross-cell-line predictions (mean RMSE=2.06 vs. 2.85). The F1-score of binary classification was 18% higher in DeepCORE than in DeepChrome (mean f1-score =0.79 vs. 0.671). These results implied that the patterns captured by DeepCORE likely represented general relationships between genetic, epigenetic, and transcriptional changes.

### DeepCORE identifies regions with biologically meaningful histone markers

We examined the distributions of HMMs in genomic regions receiving attentions in DeepCORE DNN. Our results show HMMs were present in most attended bins (**Fig. 5A**). We then randomly selected 25 genes that were transcribed above the median cutoff value and 25 genes transcribed below the median cutoff in the E071 cell line. We extracted bins with highly significant attentions and counted the presence of HMMs in the corresponding regions (**Fig. 4B**). Among genes with high transcription level, the attended genomic regions were enriched with H3K4me3 and H3K27ac that are known marks of active promoters and enhancers to enhance transcription^36,37^. Conversely, the enrichment of H3K9me3 and H3K27me3 in the attended regions near low-transcription genes were consistent with their known roles in formation of heterochromatins to repress transcription ^38^.

**Figure 4:**
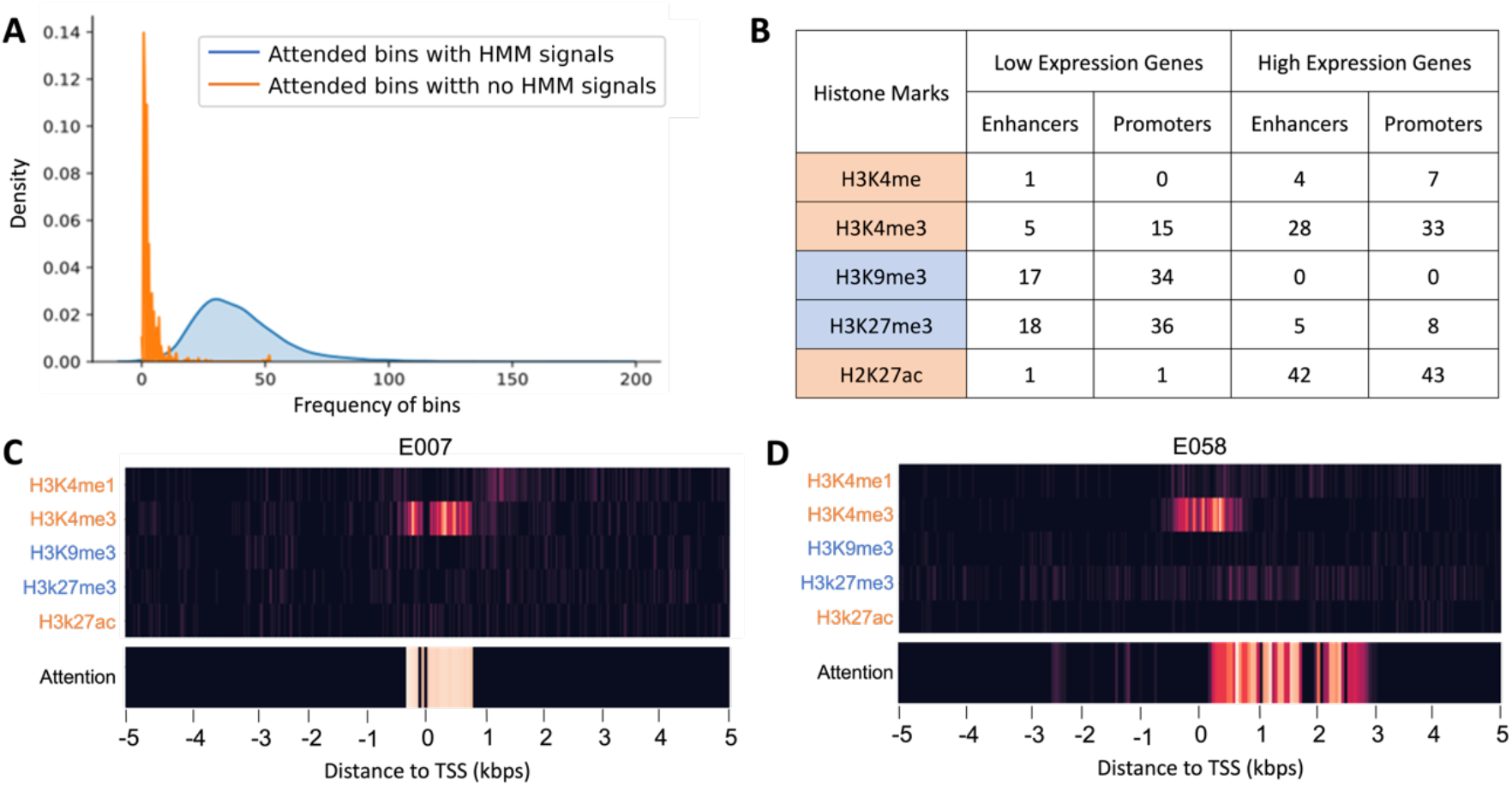
Distribution of HMMs in attended regions. (**A**) Density plots show count of attended bins with HMMs in comparison with count of attended bins with no HMMs. (**B**) Counts of bins with specific HMMs in attended regions. Data were from randomly selected 25 highly expressed genes and 25 low expression genes. Transcription activating HMMs are in orange background. Transcription repressing HMMs are in blue background. (**C, D**) Heatmaps show the raw HMM read counts and DeepCORE attention probabilities for the *CYFIP2* gene. Transcription of this gene was low in the E007 cell line (C) and high in the E058 cell line (D). The ±5kbp TSS-flanking region is encoded into 200 bins each with an attention probability.

**Figure 5:**
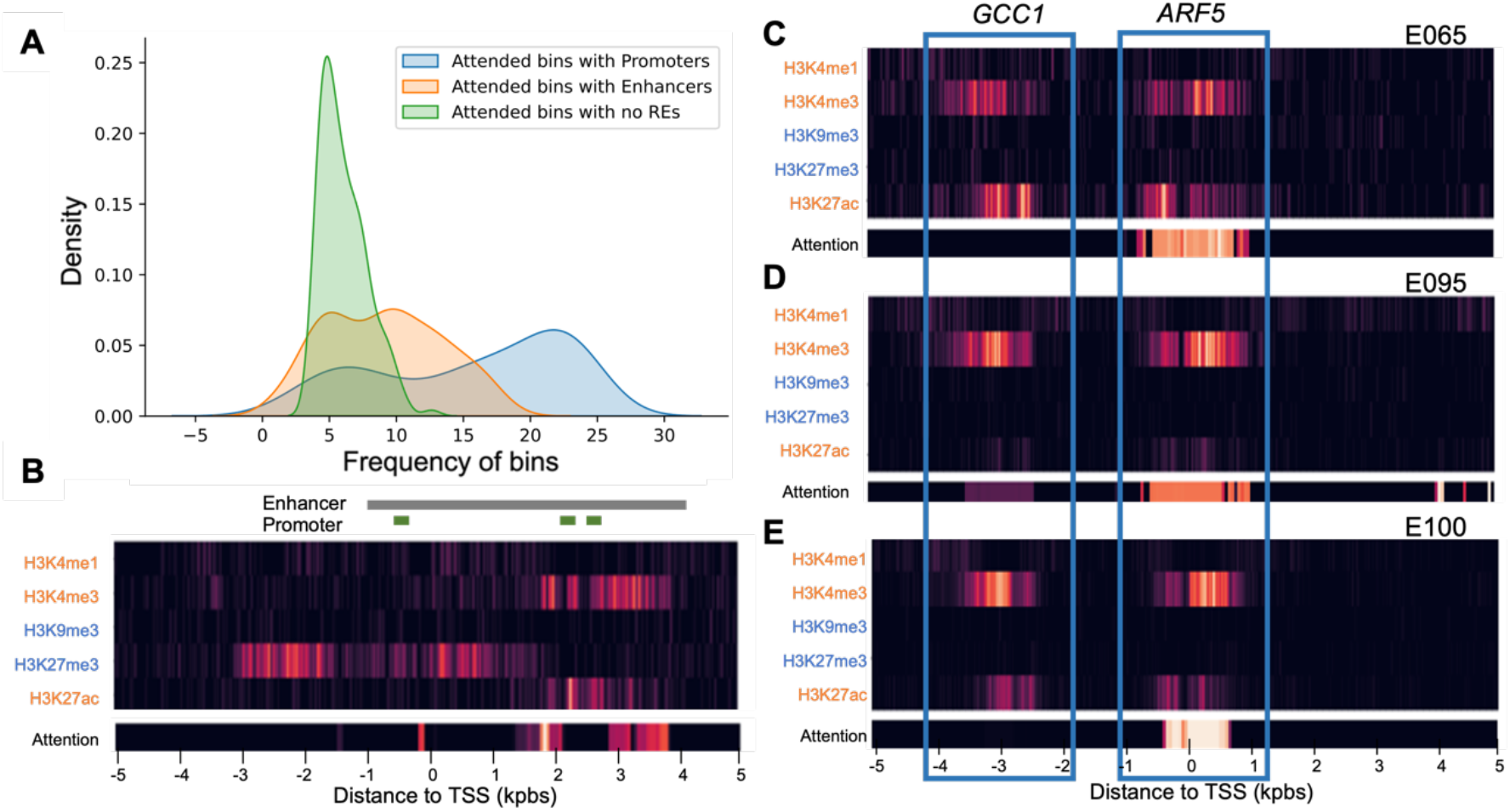
Attention analysis for regulatory elements (**A**) Density plot of the frequency of attended bins with known promoters or enhancers across 25 cell lines in comparison to random bins with high attention scores. (**B**) In the *TMEM88* gene, attended bins matched to known enhancers and promoters. HMM signals from repressing HMMs did not receive attention. (**C-E**) In the *ARF5* gene, HMM signals form two clusters (indicated with blue boxes). The right cluster mapped to the promoter of the *ARF5* received attentions. The left cluster mapped to the promoter of another gene *GCC1* did not receive attention.

Further analysis of the attended regions of the *CYFIP2* gene in two cell lines revealed interesting patterns. In the E007 cell line where *CYFIP2* gene was highly expressed, DeepCORE paid attention to regions that were close to the TSS and were occupied with the active histone mark H3K4me3 (**Fig. 4C)**. In contrast, in the E058 cell line where this gene was lowly expressed, DeepCORE paid attention to regions that were downstream of the TSS and were occupied with the repressor histone mark H3K27me3 marker and avoided regions with the activator histone mark H3K4me3 around the TSS (**Fig. 5D**). These results provided evidence that DeepCORE selects regions that are biologically relevant and reflect the underlying mechanisms of transcription regulation.

### DeepCORE can identify and fine map regulatory elements

The Eukaryotic Promoter Database (EPD)^39^ contains a comprehensive list of 29,598 experimentally validated human promoters. The GeneHancer^40^ database annotated 250,512 candidate enhancers in the human genome. We then scanned our attended regions to identify the presence of these known promoters or enhancers. To match the attended regions with the promoters, we restricted the attended regions to be within 1kbps around the TSS. No such restrictions were applied for matching enhancers.

We hypothesize that the attended regions identified by DeepCORE were enriched with known REs (i.e. promoters and enhancers annotated in EPD and GeneHancer databases. To test this hypothesis, we calculated the frequency of the attended regions containing known REs across all cell lines and the frequency of the remaining regions. On average, each gene had 23 attended bins containing known promoters and 10 attended bins containing known enhancers, but only 5 attended bins containing no known REs (Mann Whitney p-value = 4.002e^-25^ and 6.11e^-18^ respectively, **Fig. 5A**).

The *TMEM88* gene is a representative example in which the attended bins matched known promoters and enhancers. *TMEM88* is highly expressed in the E004 cell line. The ±5kb TSS-flanking region is occupied with various active and repressive HMMs. The EPD database annotated three promoters for this gene, one immediately upstream of the TSS and the other two towards the downstream. The enhancer annotated in the GeneHancer database spans a wide range starting at 1200 bps upstream of the TSS till 4000 bps downstream of the TSS. These REs all matched to the attended bins identified by DeepCORE. Furthermore, although the repressive HMM (H3K27me3) had high read counts, the DeepCORE model didn’t pay attention to it. Instead, activating HMMs received attention, which was consistent with the high transcription level of *TMEM88* in this cell line. Interestingly, inside the annotated enhancer region that spans more than 5kbps, only 30 bins covering 1,500bps received attention from DeepCORE. Because only attended bins were used to predict gene transcription level, they likely were more relevant to transcription regulation than the unattended bins covered by the enhancer.

Another interesting example is the *ARF5* gene. HMM signals in three cell lines (E095, E065, and E100) consistently highlighted two regions (**Fig. 5C-5E**). The right region corresponded to the promoter of this gene and received DeepCORE attention. The left region was 2,500bps upstream of the TSS and corresponded to the promoter of another gene *GCC1*. DeepCORE correctly identified the histone signals corresponding to *ARF5* gene and does not pay attention to the left peak. These results show strong evidence that DeepCORE identifies and fine-maps REs.

### Concordant attentions identify putative COREs

The interpreter of DeepCORE includes correlation analysis of attentions across cell lines to discover COREs. As an example, we examined the *PSMD8* gene that was consistently highly expressed across 25 different cell lines. We retrieved the attention vectors of this gene from 25 cell lines and their pair-wise correlations (**Fig. 6A**). At FDR rate <0.05, we found two blocks for which DeepCORE attentions were highly correlated (**Fig. 6B**). The first block is centered around the TSS and the second block is 3kbp downstream of the TSS. These two blocks received concordant attention across cell lines, implying that they jointly regulate transcription of the *PSMD8* gene. Indeed, these two blocks corresponded to the promoter and the enhancer of this gene. A few short regions outside the two blocks also received DeepCORE attention. However, because they did not show concordant patterns with each other, they likely are not required to function concurrently.

**Figure 6:**
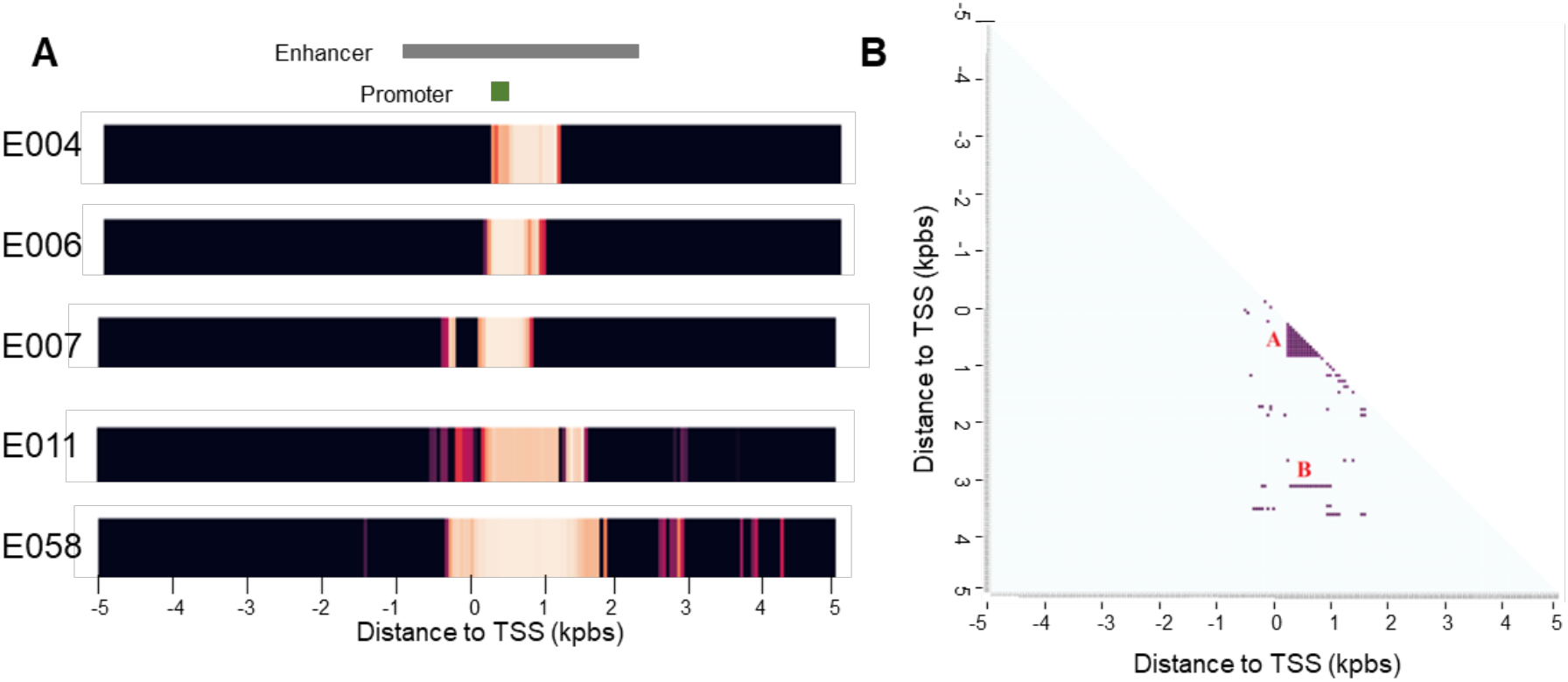
COREs in the *PSMD8* gene: (**A**) Heatmaps show attentions in 5 cell lines. (**B**) Correlation plot shows two blocks (A and B) with significant correlated attentions.

## Discussion

Multiple REs interact to regulate gene transcription. We designed the DeepCORE architecture to consider such inter-dependences in multiple aspects. At the input step, it uses two views to capture genetic and epigenetic features. At the DNN modeling step, it uses BiLSTM to allow short-range and long-range interactions, At the interpretation step, it detects correlated attentions between genomic regions. By training the DNN to predict gene transcription level based on genetic and epigenetic features within the ±5kbps TSS-flanking region, DeepCORE learns the most informative regions that are relevant to gene transcription regulation.

DeepCORE outperforms other methods on predicting gene transcription in 25 diverse cell lines. The high accuracies are strong evidence supporting that the DNN model captures informative features relevant to transcriptional regulation. It builds the foundation for subsequent analysis to further interpret the results, specifically attentions paid to each genomic region, to help mapping promoters, enhancers, and other REs. We further introduce COREs that are REs receiving concordant attentions across multiple cell lines. Using the *PSMD8* gene as an example, we showed that the COREs are a pair of promoter and enhancer that jointly regulate gene transcription.

DeepCORE uses only five HMMs as epigenetic features. However, many other types of epigenetic signals, such as DNA methylation and transcription factor binding, provide complementary information to HMM. Including these additional features may further increase the prediction accuracy and enhance the RE identification. Currently, DeepCORE examines ±5Kbps TSS-flanking region where promoters and proximal enhancers reside.

Expanding the range to 2Mbps will allow us to detect distal REs. Furthermore, as enhancers are often clustered and selective activation of different enhancers in the same cluster is tissue-specific^41–43^, concurrent modeling of multiple tissues is promising to capture the boundary between these enhancers and subsequently increase the resolution. This will also identify tissue-specific gene-promoter and gene-enhancer interactions, which is valuable knowledge that has not been annotated in existing databases.

In summary, DeeepCORE is a novel method to catalog cis-acting REs and COREs that influence gene transcription in tissue and cell-line specific context. This knowledge can be used to discover novel REs and prioritize existing REs, which will help improve our understanding of transcription regulatory mechanisms.

## Financial Disclosure Statement

This work was supported by the National Institutes of Health [grant number R01LM013438].

## Acknowledgments

The authors wish to thank all members of the Li Liu’s lab for insightful discussions on this topic.

## Data availability

The Histone Modification signals can be downloaded from https://egg2.wustl.edu/roadmap/data/byFileType/peaks/consolidated/narrowPeak/ and the gene expressions can be downloaded from https://egg2.wustl.edu/roadmap/data/byDataType/rna/expression/.

## Competing interests

The authors declare no conflict of interests.

## Author Contributions

L.L. conceived the study. L.L. and P.B.C. designed the methodology and experiments. P.B.C., H.C curated data required for the analysis and implemented the software. P.B.C., H.C., N.A., and M.L. performed analysis and visualization. P.B.C. and H.C. implemented the software. P.B.C and L.L wrote and edited the manuscript. All authors read and approved the final manuscript.

---

**Supplementary Table 1.** Different hyperparameters tested in grid search.

| Parameter | Values |
| --- | --- |
| Convolution filter size (K) | {5, 10, 50*, 100, 250} |
| Number of convolution filters ($N_{CNN}$) | {50*, 100} |
| Convolution pool size (p) | {5, 10, 50*, 100, 200} |
| Number of LSTM units ($N_{LSTM}$) | {10, 15*, 25, 50} |
| Number of attention units ($N_{ATTN}$) | {10, 25*, 50} |
| Number of fully connected network neurons ($N_{FNN}$) | {50, 100*, 200, 500} |

\* Optimal values selected after grid search

## Notes

### Competing Interest Statement

The authors have declared no competing interest.

